# Valency of Ligand-Receptor Binding from Pair Potentials

**DOI:** 10.1101/2023.09.12.557452

**Authors:** William Morton, Robert Vácha, Stefano Angioletti-Uberti

## Abstract

Molecular dynamics simulations have been crucial for investigating the dynamics of nanoparticle uptake by cell membranes via ligand-receptor interactions. Use of coarsegrained models has enabled evaluation of the effects of nanoparticle size, shape, ligand distribution on nanoparticles surface, or used thoroughly in the past decade, where a percentage of lipid heads, receptors, are attracted to sites on the nanoparticle surface, ligands. However, when pair-potentials are used to represent ligand-receptor interactions, the number of receptors interacting with one ligand, valency, may vary. We demonstrate that the curvature of a nanoparticle, strength of ligand-receptor interactions, and ligand or receptor concentration change the valency - ranging from 3.4 to 5.1 in this study. Such change in valency can create inaccurate comparisons between nanoparticles, or even result in the uptake of smaller nanoparticles than would be expected. To rectify this inconsistency we propose the adoption of a model based on bond-formation and use it to determine the extent to which previous studies may have been effected. This work recommends avoiding pair-potentials for modeling ligand-receptor interactions to ensure methodological consistency in nanoparticle studies.

**TOC Graphic:** A rendering of a ligand coated nanoparticle coming into contact with a lipid bilayer membrane. The receptor in the membrane is highlighted for clarity.

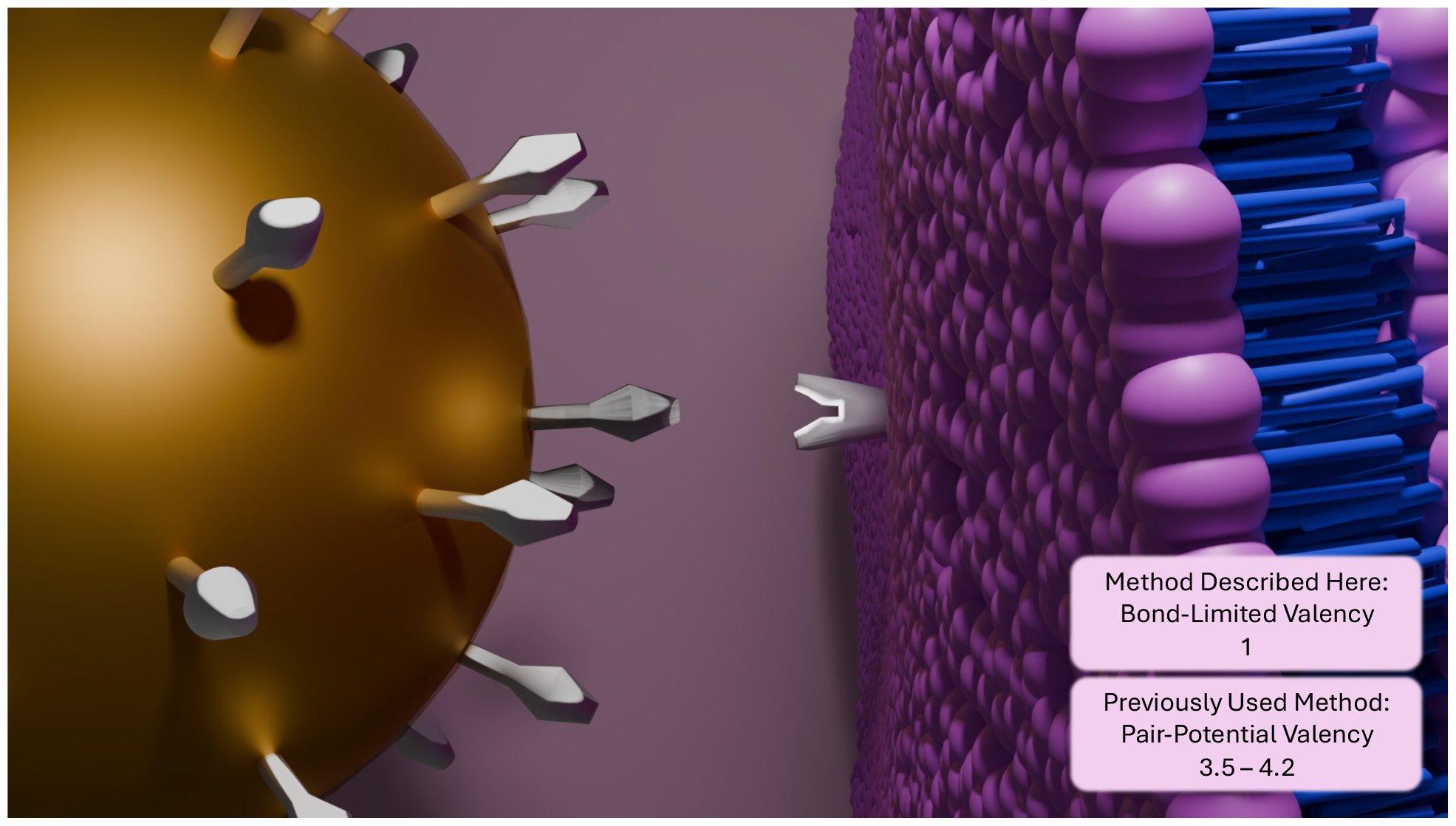

## Introduction

Over the past three decades, the use of nanoparticles as drug carriers has gained considerable attention^1,2^. The synthesis of various nanoparticle shapes has been leveraged to create nanoparticles with multiple functions, allowing them to be utilised for drug delivery, cell-imaging, and photothermal therapies^3,4^. However, targeted transport and cellular uptake still presents several scientific challenges. As an example, nanorods with an aspect ratio of 11.5 were uptaken in HeLa cells to a greater extent than those with smaller aspect ratios, but the mechanisms behind this phenomenon are not fully understood^5,6^.

Ligand-receptor valency is an essential concept in cancer research, where the population of ligands on nanoparticles could be optimized to target specific receptors on a cancer cell surface.^7^ Such optimization relies heavily on a detailed understanding of ligand’s valency.^8^ For instance, transferrin receptor 1 has been used as a ligand on the surface of nanoparticles to target cancerous cells selectively.^9,10^ Transferrin 1 is an ideal candidate because it is a signalling agent for endocytosis, often overexpressed on cancers cells, and its valency is known to be two, i.e. two ligands can bind to one receptor. ^11–13^

Lipid vesicles can be employed as simplified yet representative models of the cell membrane, due to their higher reproducibility and improved compositional control.^14^ Here, the uptake of nanoparticles is a spontaneous process — independent of cell signalling pathways, unlike, for example, micropinocytosis.^15–18^ During the wrapping process, the lipid bilayer deforms around the nanoparticle surface, resulting in an increase in bending energy. The Canham-Helfrich Hamiltonian describes the bending energy per unit surface area, as seen in Equation 1.^19^

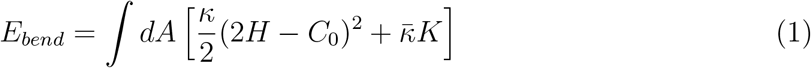

In this equation *H* and *K* are the mean and Gaussian curvature of the surface, κ and 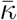 are the bending and saddle-splay modulus, respectively, and *C*_0_ is the spontaneous curvature. The increased energy that comes with wrapping (Equation 1), is balanced by the formation of ligand-receptor bonds. ^20^ As long as the energy released from the bonds is greater than the energy needed to bend the membrane, wrapping will continue.

The bending energy required to wrap a nanosphere is higher for those with a smaller radii, which have larger curvatures. With non-spherical nanoparticles, such as nanocubes, nanorods, or nanostars, this relationship can be more difficult to interpret.^21^ For example, it might be energetically favorable for the particle to change orientation while wrapping, as seen with nanorods. Therefore, analyzing how the lipid membrane wraps around the nanoparticle is essential for understanding how cells will interact with them. Unfortunately, imaging this process requires cryogenic electron microscopy. So, trial and error are needed to find the right concentration and incubation time before freezing, to properly capture nanoparticle entry; an expensive and difficult process.^22,23^

Molecular dynamics (MD) simulations of lipid membrane systems offer details about membrane wrapping, capturing the mechanisms and kinetics of nanoparticle uptake. In MD, the uptake dependence on nanoparticle shape, size, and ligand distribution can be analyzed.^24–28^ A standard computational model for these studies relies on ligand-receptor interactions, but with no specific ligand or receptor considered. Instead, the model is based on the general concept of a short-range adhesion that induces membrane wrapping. While ligands can also be used to translocate nanoparticles through the membrane, here we focus on those that reduce the membrane’s energetic barrier for wrapping a nanoparticle. *In vivo*, these short-range interactions are usually valence-limited, implying that each nanoparticle ligand and membrane receptor have a maximum number of binding partners.^29,30^ This limitation is primarily due to excluded volume effects, which prevent extra receptors from binding to the same ligand.

However, in many published articles where MD is used to model nanoparticle endocytosis, ligand-receptor interactions are described using a purely distance-dependent pairwise interaction. These interactions imply that the valence limitation is not explicitly enforced or controlled.^21,25,26,28,31,32^ Additionally, popular MD engines for coarse-grained simulations such as LAMMPS and ESPResSO lack functions to limit the number of interactions in a pair potential.^33,34^ While a ligand with a valency above 1 is not inherently inaccurate, these systems are specific, and the valency should be highly controlled within the experiment. This paper focuses on delineating the non-physical effects that can arise in this context, as well as proposing a simulation strategy to rectify them.

## Methods

### Membrane Parameterization

All simulations were performed using the LAMMPS program.^33^ Due to the substantial size of most nanoparticle structures, a coarse-grained system is necessary to investigate nanoparticle wrapping. For this reason, coarse-grained membranes have maintained their popularity over the years, and we will focus on the widely used Cooke-Deserno 3-bead lipid model. These lipids are made up of 3 beads, one bead to represent the lipid head, and the other two the lipid tail^35^. The three beads are typically connected using a finite extensible nonlinear elastic (FENE) bonds, as depicted in Equation 2, and the angle between all three beads in the lipids is maintained using the harmonic potential seen in Equation 3. The FENE potential as used in LAMMPS includes the Leonard Jones potential for a short-range repulsion.

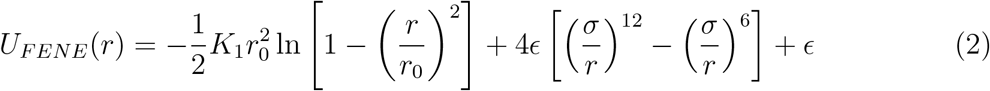

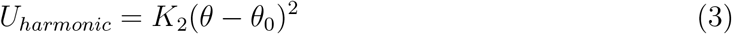

In Equations 2-5, *r* is the distance between two beads, *ϵ* is the Leonard-Jones unit for energy, and *σ* is the Leonard-Jones unit for distance. The equilibrium bond length between two beads in Equation 2, *r*_0_, is 1.5*σ*, and K_1_ is the spring constant with a value of 30*ϵ*. A harmonic potential is applied to maintain the angle formed by the three beads of each ligand in Equation 3, θ_0_, to 180^*0*^, using a spring constant, *K*_2_, of 10*ϵ*.

The pairwise potentials between individual beads are described by the Weeks-Chandler Anderson (WCA) potential in Equation 4, and the cosine potential used for lipid tails is seen in Equation 5. Equation 4 describes the non-bonding component of the potential energy between individual beads of the lipid model, which in practice, is a short-range repulsion that determines the relative size of each bead. Equation 5 facilitates the formation of a lipid bilayer by creating an attractive potential between lipid tails.

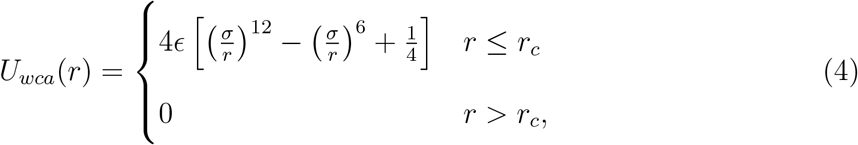

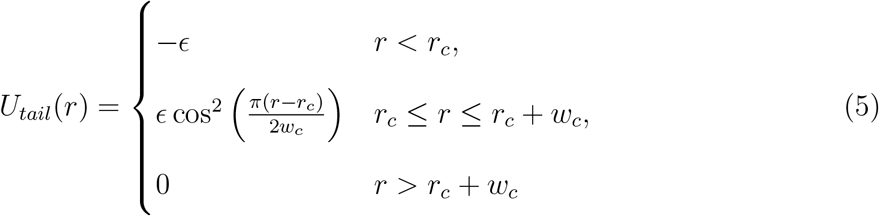

The critical length in Equation 4, *r*_*c*_, with *r*_*c*_ = 2^1*/*6^b, where *b*_*head,head*_ = *b*_*head,tail*_ = 0.95*σ* and b_*tail,tail*_ = *σ* are used to maintain a lipid membrane with no intrinsic curvature. ^36^ Altering the ‘b’ for each beads species is used to determine the relative exlcusion radius of each bead. The parameters set here are as they were originally developed by Cooke and Deserno.^37^As in Equation 2, *ϵ* is the unit of energy used to describe the potential well depth. An additional cutoff parameter (*w*_*c*_) can be altered with the system’s temperature to change membrane rigidity.

### Simulation Description

The simulated membrane used in this study is comprised of 10,452 lipids, 50% of which are receptors which can bind to ligands on the nanoparticle surface. While this results in a larger density of receptors than is biologically relevant, it decreases the simulation time by eliminating diffusion-limited interactions, and by relation the nanoparticle wrapping time^38^. There is no difference between a lipid and receptor, except an extra attractive force added between receptors and ligands. The equilibrated membrane dimension is 77 × 77 *σ* with a WCA cut-off of 1.5*σ* and k_*B*_T = 1*ϵ* maintained by a Langevin thermostat with a dampening parameter of1.0τ ^37,39,40^. A larger bilayer is used for nanospheres exceeding 12*σ* in radius, which is made of 16,296 lipids and is 98 × 98 a when equilibrated. We simulate a tensionless membrane, where the zero-tension condition is maintained using a Nose-Hoover barostat in the XY plane, parallel to the membrane, with a pressure dampening term of 1e3τ. Each system was simulated five times for 10^4^τ, with a time-step of 0.01τ.

Ligand-receptor interactions can be modelled using inter-atomic interaction potentials. A common practice is to use the pair-potential in Equation 5 that is scaled to be shorter by multiplying the WCA cut-off by 0.3 (WCA_*LR*_ = 0.45), and making the strength of the interaction *ϵ*_*LR*_ ^25^. The scaling is intended to help mimic the size and separation of ligand-receptor pairs, and here unless otherwise stated, *ϵ*_*LR*_ = 20*ϵ*. Using Equation 5 is a convenient solution, reducing the number of pair-potentials one must introduce into the system. However, for most MD engines, Equation 5 is only available as a pair-potential, where valency can not be imposed.

The nanoparticles are generated so that each bead on the surface is roughly 1a away from it’s neighbors. Each surface bead utilises the same WCA repulsive interaction between the head and tail beads of lipids to ensure they do not enter the nanoparticle volume.

The thickness of the bilayer membrane can be compared to experimental results, giving *σ*. The diffusion of lipids in the membrane can be used similarly to approximate τ. Finally, the elastic properties of the membrane can be used to find *ϵ*. To do so, we recommend the following methods within, which found that membranes similar to ours have *σ* = 0.9*nm*, and τ = 1*ns*. However, these values will be specific towards the experimental comparison being used. Since we show an effect of the parameterization here, rather than a specific experimental system, we will continue using Lennard-Jones units.

### Imposing and Measuring Valency

In the suggested method where valency is imposed, a tabulated version of the potential in Equation 5 is used as a bond style in LAMMPS. Once an unbound ligand and receptor come within 2*σ*, there is a 50% chance that a bond will be created between the two. The probability of bond formation can be altered to introduce more competition between nearby receptors, a factor which was unnecessary in this study. Bond breaking is allowed within LAMMPS, but caution must be taken to ensure that the bond energy at rupture is larger than the possible fluctuation in temperature. For this specific system, rupture is set to occur when a bond length exceeds 1.5*σ*, corresponding to a ligand-receptor binding energy of 1.25*ϵ*. Since the mean temperature is 1*ϵ*/*k*_*B*_ with a standard deviation of 4.4×10^*-*3^*ϵ*, there is no concern about a bond length exceeding the dissociation cutoff. At each time step, conditions were checked for bond formation and breaking. Therefore, since the potential is exactly the same, the only difference between the two methods is that the number of bonds that can form is fixed, maintaining a valency of 1. In this regard, we should highlight that this is not the only method available which can maintain a fixed valency. Sciortino has provided a different algorithm to effectively keep the valence fixed to 1, which introduces a repulsive 3-body potential.^41^ Whereas we only require calculations of 2-body interactions, understanding which algorithm is computationally faster, and easier to implement, is not straightforward. In fact, it likely depends on details of the system, the specifics of the MD engine used (e.g., which information is stored at each time-step) and on the exact implementation of the two algorithms.

In most coarse grained studies, the absolute value of the wrapping time is inconsequential and instead the ratio to that of a sphere is used to gauge the effect of nanoparticle size and shape. These comparisons also help to translate simulated results to experimental data.

To provide a quantitative definition of valency when using simple distance-dependent interactions, the time average ligand valency is calculated by 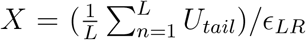 after the nanoparticle has been completely wrapped and equilibrated for 100τ. A long equilibration time ensures that the average valency is representative of the final state, where the nanoparticle is in a vesicle.

## Results and Discussion

As described using Equation 1, smaller nanoparticles should require more bending energy to wrap. A membrane must induce a large curvature to match that of the nanoparticle surface. The larger bending energy means that the total ligand receptor binding energy, found by summing the maximum binding energy per ligand on the nanoparticle surface, must increase to counter balance the bending energy.

We simulated the uptake of a range of spherical nanoparticles with radii from 7 12*σ*. In each simulation performed here, 50% of the nanoparticle surface was covered with ligands, having no imposed valency. As larger nanoparticles have more surface area, they would also have a higher number of ligands on the surface, *N*_*L*_. The interaction strength between ligands and receptors, *ϵ*_*LR*_, was scaled so that 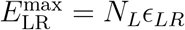, would be constant for all fully wrapped nanoparticles, regardless of their size. In scaling the interaction, we ensure that the total ligand-receptor binding energy is the same for each nanoparticle if the valency is one, isolating the effect of nanoparticle curvature on valency.

Figure 2 shows the linear relationship between the valency and Gaussian curvature of the nanoparticle. The linearity is likely due to the fractional surface area accessible to receptors in the membrane, which shrinks as a function of *R*^2^. Therefore a smaller nanoparticle means that a receptor can interact with a larger fraction of the surface. The consequence of the increased valency, is that smaller nanoparticles may have an artificially high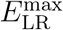, leading to an incorrect representation of the smallest nanosphere a membrane can endocytose.

**Figure 1:**
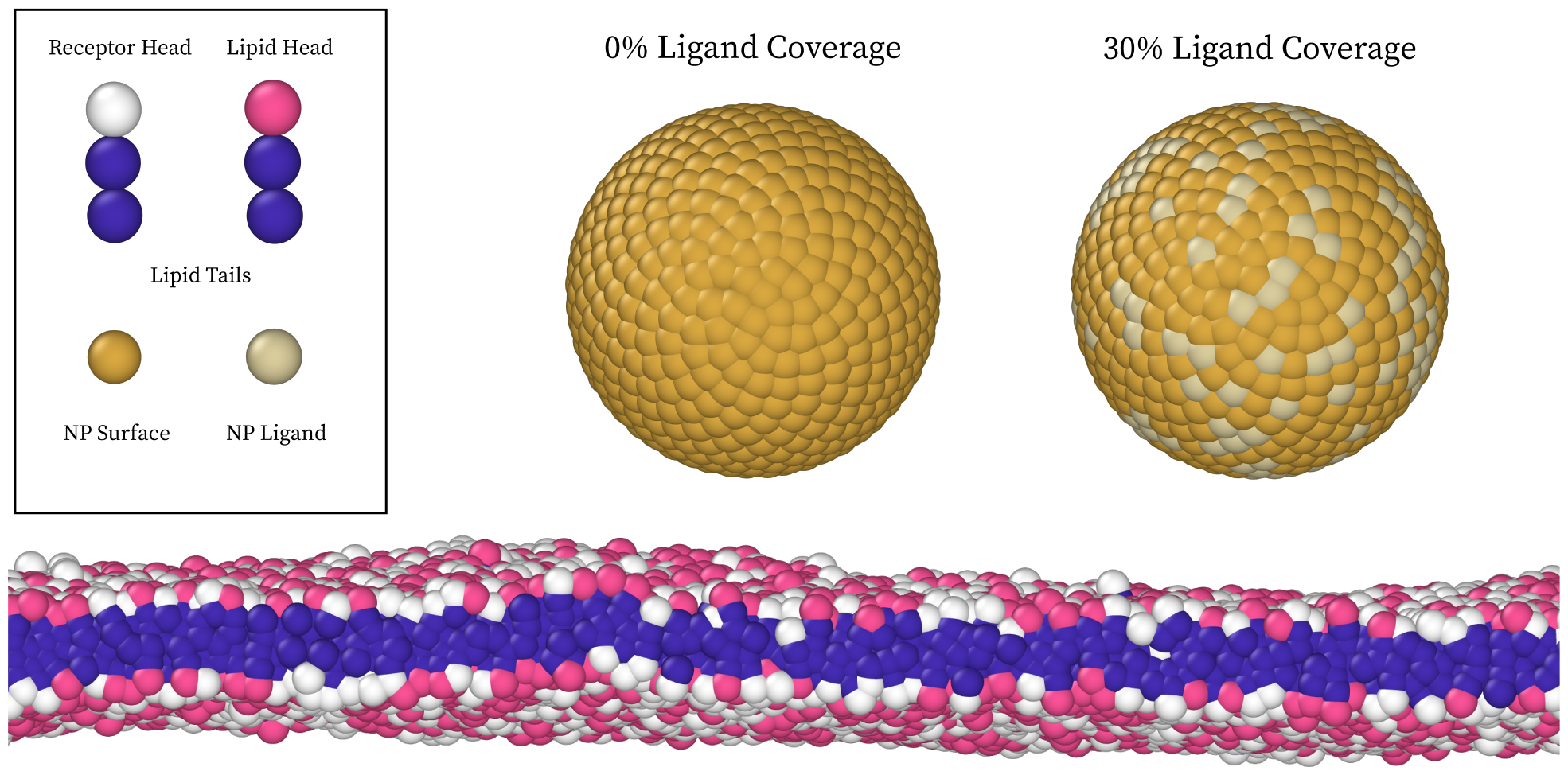
A visualisation of the lipid bilayer with two spherical nanoparticles with 0% (Left) and 30% (Right) of the surface area covered with ligands.

**Figure 2:**
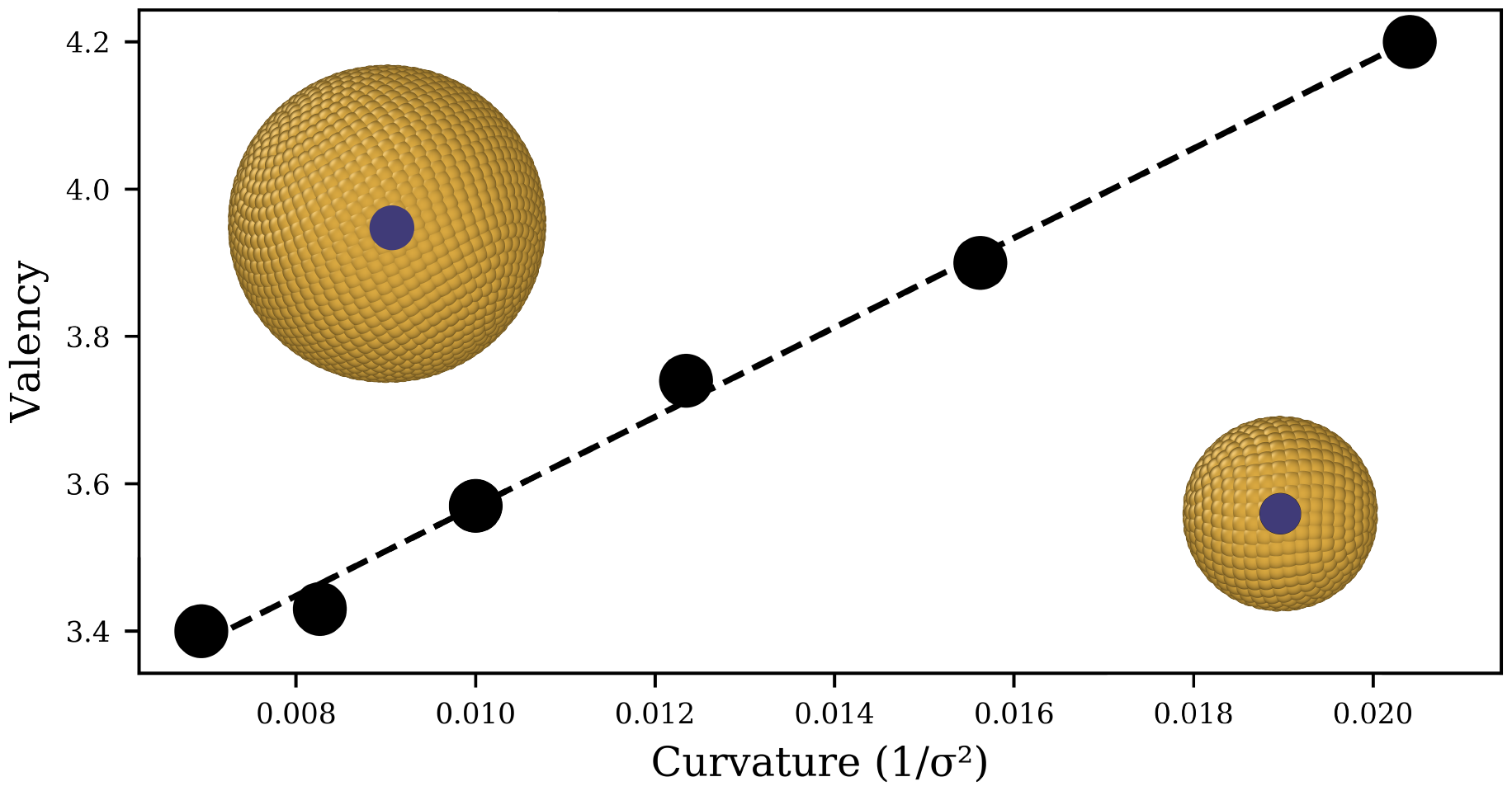
Relationship between the curvature of the nanospheres and the number of bound receptors per ligand (valency). The dotted line is a linear regression of the data points, with the coefficient of determination of 0.99. A nanoparticle of radius 12 (left) and 8 (right) *σ* are displayed, with the blue circles showing the fraction of the surface that a single lipid can interact with.

Next, the radius of the nanosphere is kept constant at 7*σ* while the ligand density and *ϵ*_*LR*_ increase. As shown in Figure 3, under these conditions the valency of ligands changes as a function of both ligand coverage and *ϵ*_*LR*_. Initially, valency increases monotonically with *ϵ*_*LR*_ at low ligand coverage. With few ligands present, receptors in the membrane are in competition to bind with ligands on the surface. A higher *ϵ*_*LR*_ causes receptors to cluster more densely, their attractive interaction with ligands counterbalancing the repulsive interaction between receptor heads. But, as ligand coverage is increased, the competitive effect diminishes, because there are more ligands to bind with, leading to a saturation effect. While this relationship is expected to hold for each nanoparticle, when valency is not regulated, the exact values are specific to the nanoparticle radius. A larger nanoparticle could accommodate more receptors, and the specific valency may change. This may also be further convoluted by the receptors ability to interact with a smaller or larger fraction of the surface area, as mentioned in Figure 2.

**Figure 3:**
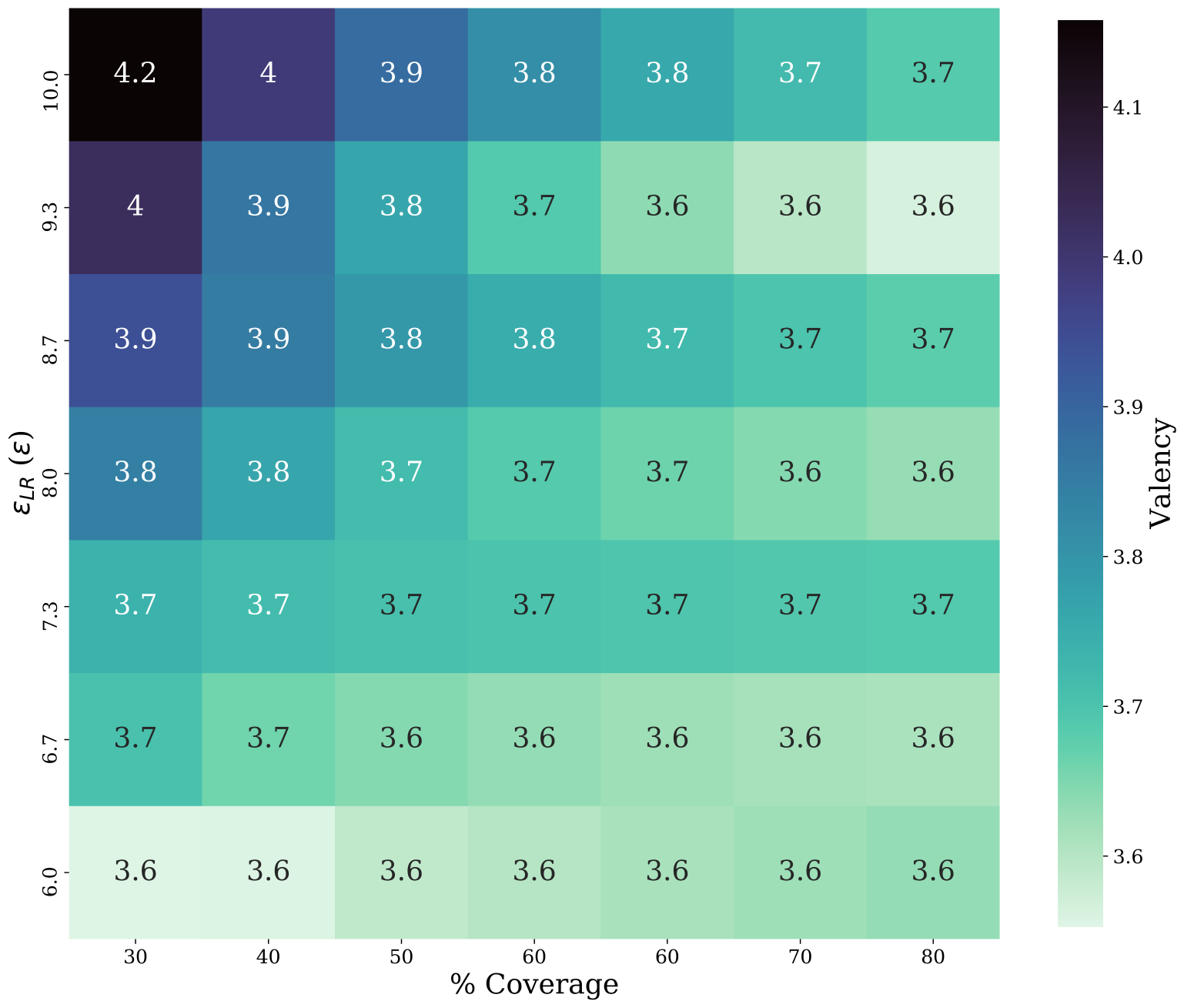
Valency as a function of ligand coverage and ligand-receptor bond energy *ϵ*_*LR*_, for a spherical nanoparticle of radius 7*σ*. Note the monotonic relationship between valency and *ϵ*_*LR*_, with a decreasing slope as ligand coverage increases.

Figure 3 suggests that simulation-based studies on nanoparticle endocytosis may contain errors, especially when not constraining valency. These errors are present when comparing effects like nanoparticle size, ligand coverage, or the strength of ligand-receptor interactions. Although a pair potential could be altered to produce a desired valency under a specific set of conditions, it would need to be tuned for each change in the system.

Finally, we test how the culmination of these effects may impact a typical study characterizing the uptake speed of a nanoparticle. Two nanospheres’ proportional uptake time should remain consistent across similar models using distance-dependent ligand-receptor interactions. Here we attempt two different approaches to modelling uptake, with and without imposing a maximum valence. Since valency is the only variable between the two systems, we can determine the scale of errors that can arise from this effect. Three membranes are used, with receptor concentrations of 20, 50, and 80% of the lipid population. As the size of the nanoparticle increases more ligands will be present on the surface, maintaining a 50% coverage. Since we are not isolating the curvature as in Figure 2, each ligand on every nanoparticle will have the same *ϵ*_*LR*_. In Figure 4, the percent error of the valence-unlimited system is shown, calculated using Equation 6

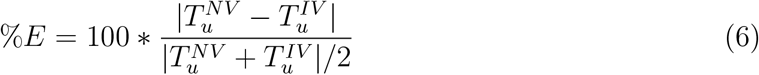

where 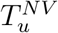 is the wrapping time of the valence unlimited system, and 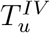 is the wrapping time of the system with an imposed valence of *X* = 1.

**Figure 4:**
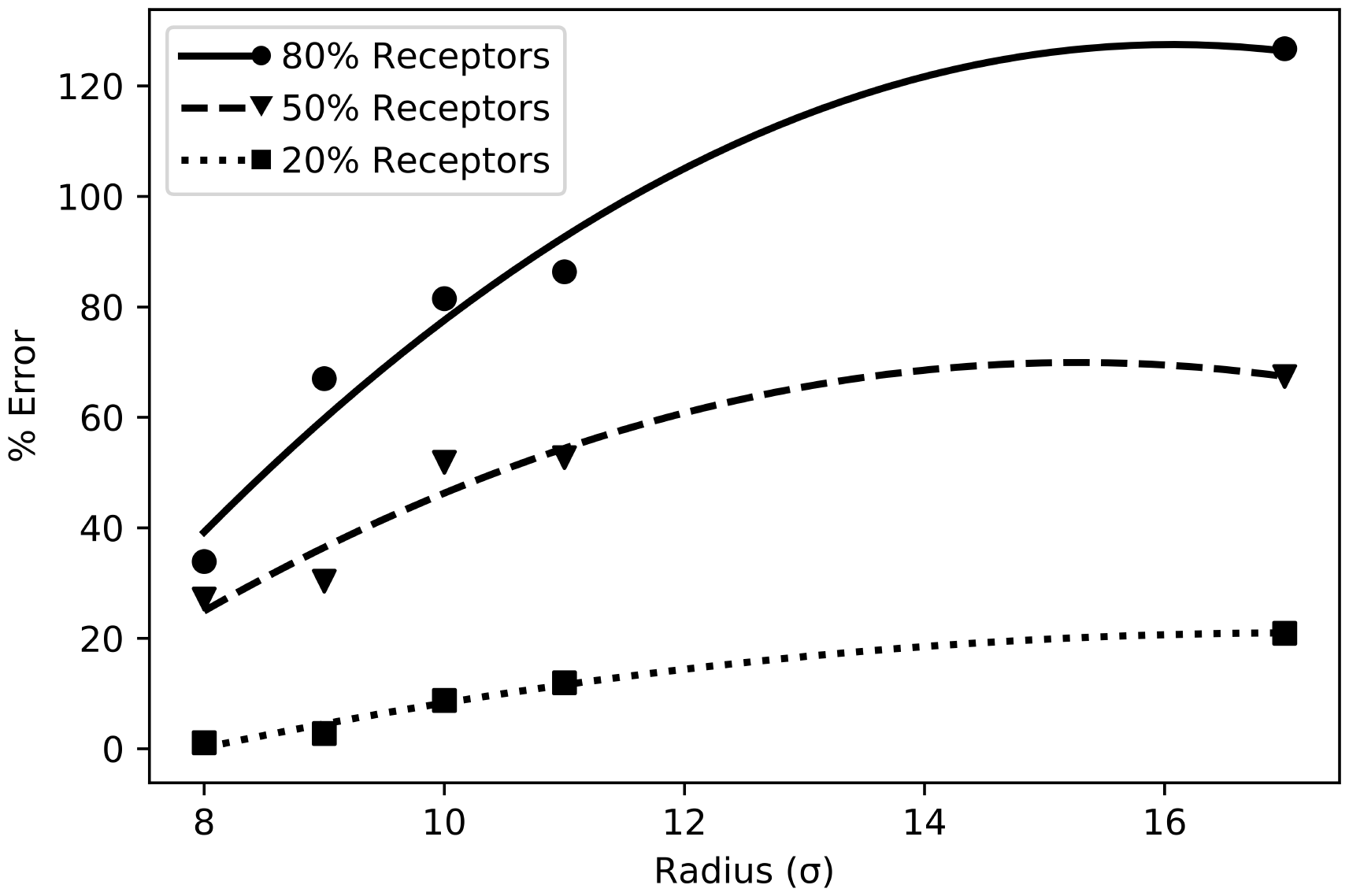
The % Error in wrapping time for the valency unlimited method of nanospheres with increasing radii, as described in Equation 6. Increasing nanoparticle radius results in a larger valency and higher errors, which is then mitigated by an insufficient number of receptors in the membrane to maintain higher valencies.

A parabolic fit is applied to the data in Figure 4 with a coefficient of determination above 0.95 for each receptor concentration, as the error is expected to increase with *R*^2^. However, a coarse grained membrane is not infinite, and using an appropriately sized membrane can reduce the need for computational resources. As a result, the membrane may have a limited number of receptors present. For example, the membrane with 20% receptors only has 1,045 receptors in total and the sphere of radius 12*σ* has 904 ligands. So, there are an insufficient number of receptors present for each ligand to have a valency of two. This suggests that membranes with a lower concentration of receptors restrict the valency in scenarios when it is not explicitly imposed. As the nanoparticle radius increases, the % Error will eventually begin to decrease as receptors become limited.

When using the same potential as a bond in LAMMPS, instead of a pair potential, it becomes easy to observe the difference in potential energy in the two systems after the nanoparticle has been endocytosed. The difference in potential energy between the two systems, divided by the potential well depth, can provide an approximate scale to the valency of the pair-potential being utilized. This assumes that all system parameters remain constant. Not every pair potential will be affected equally. However, we have used the most commonly reported one here. Therefore, we recommend that researchers utilize this method as a quick test to identify if their system has any uncontrolled valency.

Due to the artefacts presented thusfar, even scaling the effective value of the bond energy *ϵ*_*LR*_ to obtain a target valency is not a viable option. The scaling is performed by using the following equation 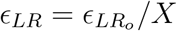, where 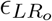 is the original ligand-receptor bond energy of 20*ϵ*. Not only does this fail to rectify the original problem, but also necessitates additional simulations to determine the scaling factors. Table 1 shows the valency for the nanoparticles in Figure 4, along with valency after *ϵ*_*LR*_ has been rescaled.

**Table 1:** Calculated valency for nanospheres interacting with a membrane made of 50% receptors. No valency is imposed, and it is shown here that the valency is not easily controlled by scaling *ϵ*_*LR*_.

| Radius ($\sigma$) | $\epsilon_{LR_o}(\epsilon)$ | $X_{Original}$ | $\epsilon_{LR}(\epsilon)$ | $X_{Scaled}$ |
| --- | --- | --- | --- | --- |
| 7 | 20 | 5.15 | 3.87 | 0.57 |
| 8 | 20 | 5.07 | 3.93 | 0.56 |
| 9 | 20 | 4.92 | 4.06 | 0.58 |
| 10 | 20 | 4.84 | 4.12 | 0.60 |
| 11 | 20 | 4.74 | 4.21 | 0.63 |
| ... | ... | ... | ... | ... |
| 17 | 20 | 4.35 | 4.59 | 0.69 |

While the error is reduced, there is still no control over the ligand valency.

## Conclusions

Variables that may be considered during a MD study on passive nanoparticle endocytosis, such as the number of ligands on a nanoparticle surface, ligand-receptor binding energy, and size of the nanoparticle, each change the valency of ligand-receptor interactions. Here, valency acts as a confounder which can misrepresent both the relationship between nanoparticles and lipid membranes, and the comparisons between nanoparticles. Attempts to mitigate the inconsistent valency by scaling the interactions between ligands and receptors can improve the accuracy of results, but not eliminate the core problem. The study suggests that not all published results will be effected equally. Studies that use lower receptor concentrations reduce the valency when not imposed due to the lower probability of multiple receptors diffusing to the same ligand. The trade-off is that lower receptor concentrations result in longer simulation times, which is unnecessary for comparing nanoparticle *T*_*u*_ if a valence-limited method is used. Those that have scaled interactions may also be able to mitigate the uncontrolled valency to a certain extent. However, the effectiveness of this method is only shown here for spheres. Given the inconsistencies in valency and wrapping time demonstrated throughout this work, the authors recommend that isotropic pairwise functions to model ligand-receptor interactions be discontinued when possible, and valency be reported in the methods of each publication.

All code necessary to recreate the results of this paper, and perform simulations in LAMMPS can be found at: https://github.com/shakespearemorton/Nano_Wrap

These simulation were performed at the Imperial College Research Computing Service (see DOI: 10.14469/hpc/2232).

