## Supplementary_Information for "Valency of Ligand-Receptor Binding from Pair Potentials"

### Valency of Ligand-Receptor Binding from Pair Potentials: Supplementary Information

#### Effect of Changing the Pair Potential

Here we demonstrate that the uncontrolled valency effects outlined in the manuscript are not an artefact of the chosen pair-potential. In Figure S1A we have graphed each of the pairwise interactions that describe the lipid interactions, as outlined in our manuscript. In short, there is a repulsive interaction between the lipid heads (HEAD_HEAD). The attractive interaction, indicated by a negative potential energy, between the lipid tails keeps the membrane intact (TAIL_TAIL).[1] The interaction between the beads making up the nanoparticle surface, and the lipid heads/tails, also takes the repulsive form of lipid head interactions (HEAD_HEAD). This repulsive interaction creates the nanoparticle surface and stops lipids from entering the structure. These pair potentials should not change in this system, as they are the building blocks that have been well verified throughout several publications.[2–4]

The form of the pair potential between a ligand (on the nanoparticle surface) and receptor (located in the membrane as a lipid head) is based on the free energy between a ligand and receptor, an example of which is seen in Figure S1B. The cosine-squared pair potential used within our simulation is plotted in Figure S1C (LIGAND_RECEPTOR). Modifications to our pair potential are limited by the system we are trying to simulate. Below we report a physical explanation of the modifications, along with graphs of each potential, and results from simulations that were performed using each method with a nanoparticle of radius = 7 $\sigma$, and 80% of the surface made up of ligands. The simulation set up is equivalent to that shown within the manuscript. An example of the ligand-receptor interaction energy is shown in Figure S1D as the nanoparticle is wrapped by the membrane.

#### Shift the potential (Figure S1E)

Shifting the potential towards smaller separation distances will introduce competition between the repulsive nanoparticle surface, and the attractive ligand-receptor interactions. As the separation distance decreases, the overall interaction between the ligand and receptors will decrease, and eventually there may be insufficient interaction energy to counterbalance the curvature of the membrane.

Shifting the minima towards larger separation distances causes defects in the membrane to occur (Figure S1H). This is because the center of the attractive interaction is away from the nanoparticle surface, and begins to extend into the vesicle bilayer. The membrane, rather than returning to a low energy flat plane, curves to maintain interaction with the ligands.

#### Expand the potential (Figure S1F)

Expanding the potential, by adjusting the Weeks-Chandler-Anderson cut-off ($WCA_{\text{cut}}$), will increase the area where receptors can bind around the ligand, resulting in a higher valency. The large attractive region allows ligands to interact with receptors outside of the vesicle, causing the membrane to change conformation to maintain contact with the nanoparticle after uptake (Figure S1H). This is a non-physical effect that is unwanted in the system.

#### Change the depth of the potential (Figure 3)

Increasing or decreasing the depth of the potential modulates the strength of the attraction between ligands and receptors, which is addressed in the manuscript (Figure 3).

#### Change the form of the potential (Figure S1G)

Several pair potentials could be used to represent the ligand receptor interactions. Here we choose to match one of the forms, although not necessarily the exact values, derived from experiments in Figure S1B using the following Equation S1.

$$V(r) = \epsilon \left[ \left( \frac{\sigma}{r} \right)^{12} - \left( \frac{\sigma}{r} \right)^{6} \right] + A \exp \left( - \frac{(r - B)^2}{2C^2} \right) \tag{S1}$$

The potential energy is a function of the separation distance, r. The form is a combination of the Lennard-Jones potential, with $\sigma$ equal to 1 and $\epsilon = 20$, and a Gaussian term, the height of which can be modulated by changing A, the center of which is at B, and the width of which is determined by C. Here A, B, and C are 3, 1.2, and 0.1 respectively. Even with the modified form, we can see that there is still an issue with valency present.

The valency from each of these experiments can be seen in Table S1:

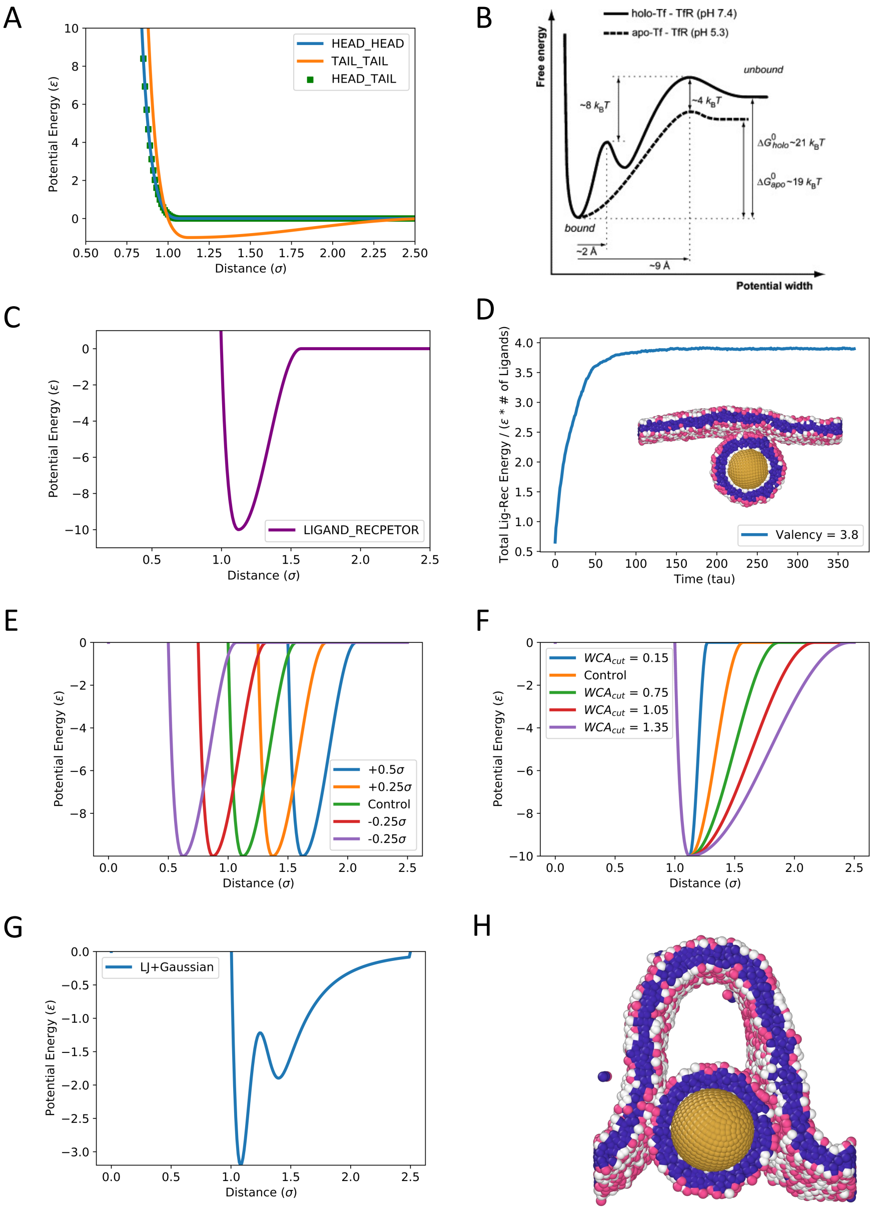


Figure S1: A) The form of pair-potentials necessary to form a lipid bilayer membrane. B) Experimentally derived interaction energy between transferrin and its receptor. Reproduced with permission from,[5] Elsevier 2024. C) The pair-wise interaction between ligands and receptors. This is based on the cosine-squared potential, as used in the manuscript. D) The total energy in the system from ligand-receptor bonds, divided by the maximum interaction energy times the number of ligands - a good approximator for the valency of the system. Inset shows the final snapshot of the simulation, with the nanoparticle inside of a vesicle. E) Modified ligand-receptor pair potentials to examine the effect of shifted potentials. F) Modified ligand-receptor pair potentials to examine the effect of expanded potentials. G) Modified ligand-receptor pair potential built from combining a Lennard-Jones potential with a Gaussian curve. The shape is intended to mimic that seen in Figure 1B. H) Buckled membrane, which is the effect of having a potential that can penetrate through the vesicle that surrounds the nanoparticle.

Table S1: Valency of modified pair potentials representing ligand-receptor interactions.

| **Method** | **Alteration** | **Graph of Pair-Potential** | **Valency** |
|---|---|---|---|
| Control | None ($WCA_{cut} = 0.45$) | Fig 1C | 3.8 |
| Shift Minima | $-0.5\sigma$ | Fig 1E | 3.2 |
| Shift Minima | $-0.25\sigma$ | Fig 1E | 3.6 |
| Shift Minima | $+0.25\sigma$ | Fig 1E | 4.6 |
| Shift Minima | $+0.5\sigma$ | Fig 1E | 5.1 |
| Expand | $WCA_{cut} = 0.15$ | Fig 1F | 3.6 |
| Expand | $WCA_{cut} = 0.75$ | Fig 1F | 5.1 |
| Expand | $WCA_{cut} = 1.05$ | Fig 1F | 7.0 |
| Expand | $WCA_{cut} = 1.35$ | Fig 1F | 9.3 |
| LJ + Gaussian | New pair-potential | Fig 1G | 4.6 |
